# Segmentation of Tissues and Proliferating Cells in Light-Sheet Microscopy Images using Convolutional Neural Networks

**DOI:** 10.1101/2021.03.08.434453

**Authors:** Lucas D. Lo Vercio, Rebecca M. Green, Samuel Robertson, Si Han Guo, Andreas Dauter, Marta Marchini, Marta Vidal-García, Xiang Zhao, Ralph S. Marcucio, Benedikt Hallgrímsson, Nils D. Forkert

## Abstract

**Background and Objective:** A variety of genetic mutations are known to affect cell proliferation and apoptosis during organism development, leading to structural birth defects such as facial clefting. Yet, the mechanisms how these alterations influence the development of the face remain unclear. Cell proliferation and its relation to shape variation can be studied in high detail using Light-Sheet Microscopy (LSM) imaging across a range of developmental time points. However, the large number of LSM images captured at cellular resolution precludes manual analysis. Thus, the aim of this work was to develop and evaluate automatic methods to segment tissues and proliferating cells in these images in an accurate and efficient way.

**Methods:** We developed, trained, and evaluated convolutional neural networks (CNNs) for segmenting tissues, cells, and specifically proliferating cells in LSM datasets. We compared the automatically extracted tissue and cell annotations to corresponding manual segmentations for three specific applications: (i) tissue segmentation (neural ectoderm and mesenchyme) in nuclear-stained LSM images, (ii) cell segmentation in nuclear-stained LSM images, and (iii) segmentation of proliferating cells in Phospho-Histone H3 (PHH3)-stained LSM images.

**Results:** The automatic CNN-based tissue segmentation method achieved a macro-average F-score of 0.84 compared to a macro-average F-score of 0.89 comparing corresponding manual segmentations from two observers. The automatic cell segmentation method in nuclear-stained LSM images achieved an F-score of 0.57, while comparing the manual segmentations resulted in an F-score of 0.39. Finally, the automatic segmentation method of proliferating cells in the PHH3-stained LSM datasets achieved an F-score of 0.56 for the automated method, while comparing the manual segmentations resulted in an F-score of 0.45.

**Conclusions:** The proposed automatic CNN-based framework for tissue and cell segmentation leads to results comparable to the inter-observer agreement, accelerating the LSM image analysis. The trained CNN models can also be applied for shape or morphological analysis of embryos, and more generally in other areas of cell biology.

## 1. Introduction

Structural birth defects and congenital anomalies are a major human health issue, accounting for 300,000 worldwide deaths each year and a significant proportion of the global burden of disease [1]. Most congenital anomalies have a genetic cause that results in a disease by perturbing cellular or molecular processes during embryonic growth and development. Interventions aiming at treating or preventing such diseases require a mechanistic understanding of these disruptions. Many publications report small changes in cell proliferation or apoptosis in response to a perturbation in a model system. However, these changes are often measured by manual or semi-automatic cell counting in a few stained tissue regions [2, 3, 4]. A major limitation of this approach is that it is difficult to investigate changes across a whole region of tissue or how these changes contribute to the overall anatomical development.

During embryogenesis, tissue layers (ectoderm, mesoderm, neural crest and endoderm) form all organs and structures of the body. Each tissue layer has different properties and specific cell fates. In this work, we specially focus on the growth and development of the face. During early development, the neural crest is actively migrating and proliferating in response to cues from the ectoderm. The neural crest mixes with the mesoderm to form a tissue type commonly known as mesenchyme, which continues to form the bones and muscles of the face. Proliferation and migration of the mesenchyme are thought to be the primary drivers of facial morphology [5, 6].

For the study of embryogenesis, fluorescence imaging allows the observation of certain biological processes by specifically displaying the molecules and structures involved. Light-Sheet microscopy (LSM), a rather novel fluorescence imaging technique, allows 3D imaging of whole biological samples, even at early developmental stages, with a spatial resolution that can display single cells [7]. Furthermore, LSM imaging can be acquired without physical slicing of the samples, thus, preserving the shape of the sample. Technically, LSM illuminates a plane in the sample using a determined frequency, whereas the fluorecence is imaged using an sCMOS camera perpendicular to the plane. The 3D image is obtained by moving the illumination plane along the sample [8]. LSM allows imaging of individual cell nuclei that have been stained for various markers. In this work, we specifically focus on the DAPI (4’,6-diamidino-2-phenylindole) marker that stains all nuclei, as well as PHH3 (phospho-Histone H3) that only stains actively dividing or proliferating cells.

Light-Sheet imaging has practical drawbacks for its analysis. The size of a multi-channel image at 5x zoom can easily approach 300 GB and include thousands of 2D images. These images are also prone to noise and loss of signal due to sample preparation variability and imaging artifacts. Due to the large number of images acquired by LSM, the large size of these images, and the different artifacts that can be present, large-scale manual analysis of LSM images is not feasible. The aim of this work was to develop and evaluate an automatic framework for segmentation of the mice embryo morphology, its tissues, cell nuclei, and proliferating cells to support developmental biology research, particularly the analysis of cell proliferation and shape change during embryo growth, and the effect of gene mutations on them.

### 1.1. Related work

In recent years, a variety of automatic methods have been applied to fluorescence microscopy, and particularly LSM, for quality improvement and analysis. The high variability of intensities across and between samples in fluorescence microscopy complicates the development of general methods for automatic analysis. This variability is a result, for example, of sample preparation, such as excessive non-uniform tissue clearing and/or antibody penetration, noise, loss of signal as the light-sheet travels through the specimen, and due to the non-specific fluorescent signal (background). To date, multiple mechanical and computational methods have been proposed to correct for these problems. Generally, mechanical methods aim to improve the LSM acquisition technique. For example, Turaga and Holy [9] proposed a method to correct for defocus aberrations by tilting the angle of the light-sheet microscopy by a few degrees, whereas the remaining aberrations can be corrected using adaptive optics. Bourgenot et al. [10] examined how aberrations can occur in single plane illumination microscopy (SPIM) of zebrafish samples, and proposed a wavefront correction method for this artefact. Among the computational methods for post-processing, Yang et al. [11] developed a method based on Convolutional Neural Networks (CNNs) to automatically assess the focus level quality in microscopy images. Yayon et al. [12] proposed a semiautomatic method to normalize images, taking into account the background intensity and signal elements while Weigert et al. [13] proposed a normalization method based on percentiles to overcome the problem of extreme signal values.

Once the image correction is done, the image quantification process (e.g. tissue segmentation and cell counting) can be performed with improved accuracy. Particularly for cell segmentation in fluorescence images, Ilastik [14] is a widely used software tool. It provides a trainable pixel segmentation tool, where the user specifies the image features to be extracted and can configure the hyperparameters of the random forest classifier used in the background. Additionally, a variety of methods based on CNNs have been proposed in recent years. For example, Ho et al. [15] proposed a 3D CNN model for segmenting nuclei in rat kidneys labelled with Hoechst 33342. The authors specifically emphasized on the large amount of annotated data required to train CNNs. To overcome this problem, they trained the CNN with synthetic data and tested it in real images. Falk et al. [16] segmented a variety of microscopy images using a generic 2D CNN for segmentation, called U-net. This CNN model was trained using datasets provided by the ISBI Cell Tracking Challenge 2015 [17] and by applying data augmentation techniques to overcome the problem of limited data for CNN training. However, to the best of our knowledge, there is no comprehensive, publicly available software solution so far that allows segmenting different tissue and cell types in LSM datasets.

### 1.2. Contribution of this Work

In recent years, U-nets have been successfully used for many medical image segmentation tasks [18]. The original U-net architecture was proposed to segment neuronal structures on 2D electron microscopy images [19]. Since then, this CNN model has been extended to segment other types of structures and data, such as the prostate [20] and ischemic strokes [21] in 3D MRI volumes. In this work, optimized U-nets are used to solve the challenging segmentation problems in 3D LSM datasets.

The segmentation of mesenchyme and neural ectoderm is a key first step to properly analyze the shapes of these rapidly growing regions at different developmental stages. Segmentation of both tissues also allows a more robust registration of images from different samples acquired at the same age and across different ages. After registration, shape changes can be studied using, for example, geometric morphometrics [22]. In this work, we train and evaluate an U-net model using a unique database of DAPI-stained LSM images with corresponding manual segmentations to automatically segment the mesenchyme and neural ectoderm.

The second key aim is to quantify cell proliferation in the mesenchyme to support basic science and clinical research investigating how mitosis and migration drive morphological changes. For nuclei as well as segmentation of proliferating cells, we re-trained the cell segmentation U-net CNN model described by Falk et al. [16] using our own LSM datasets. More precisely, we trained this CNN model using datasets of DAPI-stained images with corresponding annotated cells for nuclei segmentation. For segmentation of proliferating cells, the U-net was trained using PHH3-stained images with corresponding manual segmentations. Finally, the three segmentations are combined to create maps of relative proliferation in the mesenchyme. The source code, software, and annotated datasets have been made publicly available at https://github.com/lucaslovercio/LSMprocessing.

## 2. Materials and Methods

### 2.1. Image acquisition

Five E9.5 and five E10.5 mice embryos were harvested and fixed overnight in 4% paraformaldehyde. After fixation, they were processed for clearing and staining. The clearing step followed the CUBIC protocol [23]. Briefly described, embryos were incubated overnight in Cubic1/H20 at room temperature followed by incubation in Cubic 1 at 37^*◦*^C degrees until clear (1-3 days). Samples were blocked in 5% goat serum, 5% dimethyl sulfloxide, 0.1% sodium azide, and phosphate buffered saline at 37°C for 1.5 days. After this, they were incubated in primary antibody (Abcam ab10543) (1:250) and DAPI (1:4000) in 1% dimethyl sulfloxide, 1% goat serum 0.1% sodium azide, and phosphate-buffered saline (PBS) for 7 days at 37°C with shaking. We performed several washes in PBS for 3 days at room temperature. Next, the samples were incubated with secondary antibody (1:500) (Abcam ab150167) for 5 days at 37°C with gentle shaking. The samples were then washed for several days in PBS at room temperature and were embedded in 1.5% low melt agarose, incubated in 30% sucrose for 1 hour and then placed in Cubic 2 overnight before imaging. The samples were imaged using a Lightsheet Z1 scanner (Zeiss). Images were acquired using single side illumination at 5x zoom using a minimum of 3 laser channels: 405-DAPI, 488-background, and 647-PHH3.

### 2.2. Datasets

From the DAPI-stained scans of ten available mice embryos, random images were selected from these z-stacks for manual tissue segmentation. Particularly, 86 were used for CNN training, 36 for validation, and 54 for testing, following the usual proportion of 50-20-30% for machine learning datasets (DAPI-Tissue dataset, Table 1) [24]. These 176 images were cropped to 1024 × 1024 because the size of the images vary between the scans, depending on the size of the embryo (Fig. 1a). This patch size was determined to be feasible for human annotation, as the observers required up to five minutes segmenting each slice. Prior to further processing and manual annotation, the images were intensity-normalized using percentile-based equalization [13]. Five expert observers manually segmented the mesenchyme and neural ectoderm tissues in disjoint subsets of the DAPI-Tissue set (Fig. 1b). Each slice used for testing was independently segmented by two different observers to assess the inter-observer variability.

**Table 1:** Distribution of the images for training (train), validation (val) and test, sizes and annotated classes for the three proposed datasets.

| Dataset | Train | Val | Test | Size | Classes |
| --- | --- | --- | --- | --- | --- |
| DAPI-Tissue | 86 | 36 | 54 | $1024 \times 1024$ | Mesenchyme - Neural ectoderm - Background |
| DAPI-Cells | 96 | 17 | 55 | $131 \times 131$ | Mesenchyme cell - Neural ectoderm cell - No cell |
| PHH3-Cells | 86 | 20 | 49 | $1024 \times 1024$ | Proliferating cell - No proliferating cell |

**Figure 1:**
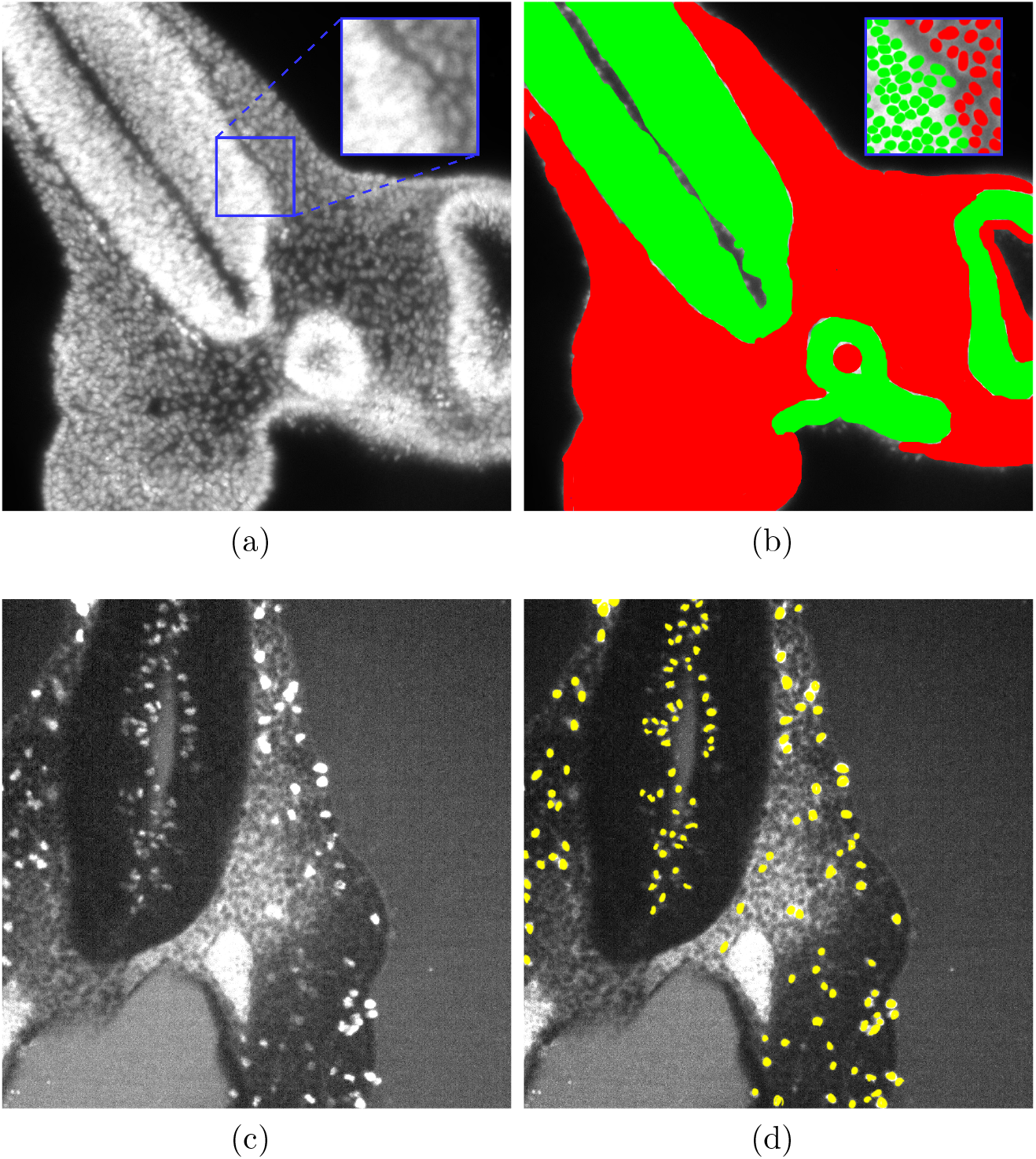
Datasets used for model development and evaluation. (a) Cropped and normalized DAPI-stained image for mesenchyme and neural ectoderm segmentation (DAPI-Tissue). In blue, subimage used for cell segmentation (DAPI-Cells). (b) Manual segmentation of mesenchyme (red) and neural ectoderm (green) of (a). (c) Cropped and normalized PHH3-stained image (PHH3-Cells). (d) Manual segmentation of proliferating cells marked (yellow).

For development of the cell segmentation model, a random sub-image with a size of 131 × 131 was extracted for manual cell segmentation from each image in the DAPI-Tissue set (Fig. 1a). Here, the observers segmented single cells, while differentiating them according to the tissue (mesenchyme or neural ectoderm) that they belong to (Fig. 1b). Heavily blurred patches, where single cells could not be separated, were removed from the dataset, resulting in a total of 168 images, whereas 96 were used for training, 17 for validation, and 55 for testing (DAPI-Cells dataset, Table 1). The latter subset was independently segmented by two different observers for variability assessment. It is worth noting that due to not all patches containing both types of cells, the proportion of images for training was increased in this case. Finally, for the development and evaluation of the CNN model for segmentation of proliferating cells (PHH3-Cells), 155 images were randomly selected from the PHH3 scans of the ten embryos, whereas 86 images were used for training, 20 for validation, and 49 for testing. These images were cropped to 1024 × 1024 and intensity normalized [13] (Fig. 1c). In this case, the observers segmented proliferating cells (Fig. 1d). The patch size used for this task was different from the images used for cell segmentation in the DAPI-stained scans because the proliferating cells represent only a small fraction of the total number of cells. Thus, a human observer can segment a larger image in a similar time frame. Similarly to the previous two subsets, these test images were independently segmented by two different observers for variability assessment.

### 2.3. LSM segmentation workflow

Fig. 2 shows the proposed workflow to generate the map of proliferating cells in the mesenchyme of an embryo. The two inputs are the channels belonging to DAPI and PHH3 in light-sheet microscopy scans, and the two main outputs are the segmented mesenchyme and neural ectoderm tissues and the relative proliferating cell volume for mesenchyme. The methods used for tissue segmentation, cell segmentation, and proliferating cell segmentation are detailed in the following sections.

**Figure 2:**
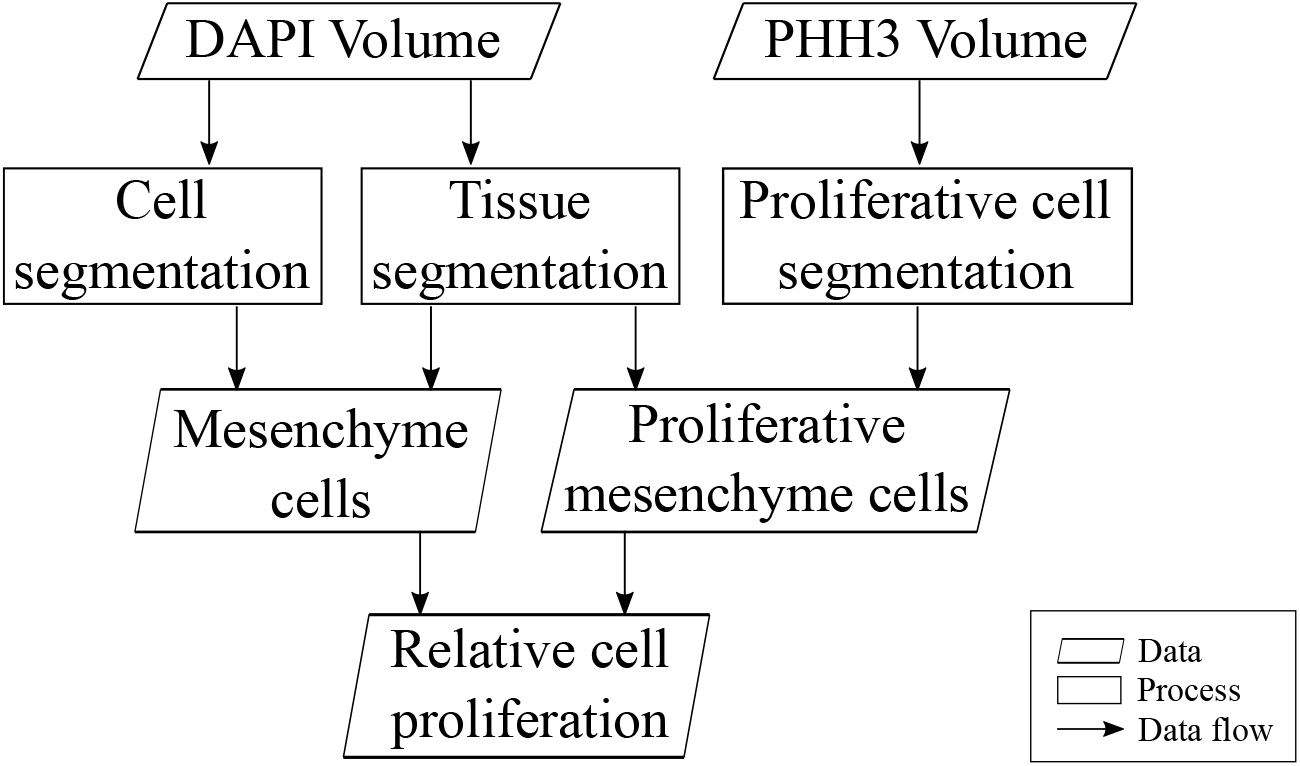
Overview of the proposed light-sheet microscopy segmentation workflow.

### 2.4. Tissue segmentation

For the task of segmenting the mesenchyme and neural ectoderm in DAPI-stained images, and distinguishing these tissues from the background, a simple U-net CNN model was developed (Fig. 3). Therefore, a large random search was conducted over various settings of hyper-parameters using the DAPI-Tissue training and validation sets. The hyper-parameters investigated included the batch size, learning rate, number of filters, kernel size, and optimizer method. The search began with a high-dimensional space containing a wide range of hyper-parameters and a wide range of possible settings for each hyper-parameter. At each stage, a large number of models with randomly selected hyper-parameter configurations (within predefined ranges) were trained, and the settings of the best models were recorded. From a manual examination of the settings with the best results, the ranges of possible hyper-parameters were refined after each stage. After the space was reduced to a feasible size, a grid search was conducted over the remaining possible hyper-parameter values. Finally, the best model was trained and validated five times to ensure that its performance was not due to random factors inherent in the training process such as the initialization of the weights. Using this procedure, the best model configuration identified used a batch size of 8, an equally weighted combined loss function of soft-Dice and categorical cross-entropy, batch normalization, and ReLU activations in the hidden layers [25, 26] (Fig. 3). This model was trained using the RMSprop optimizer with a learning rate of 1E-4 for 200 iterations. Training data augmentation (flipping, blurring, affine deformations) was performed on the fly [27] with two dropout layers at the end of the contracting path.

**Figure 3:**
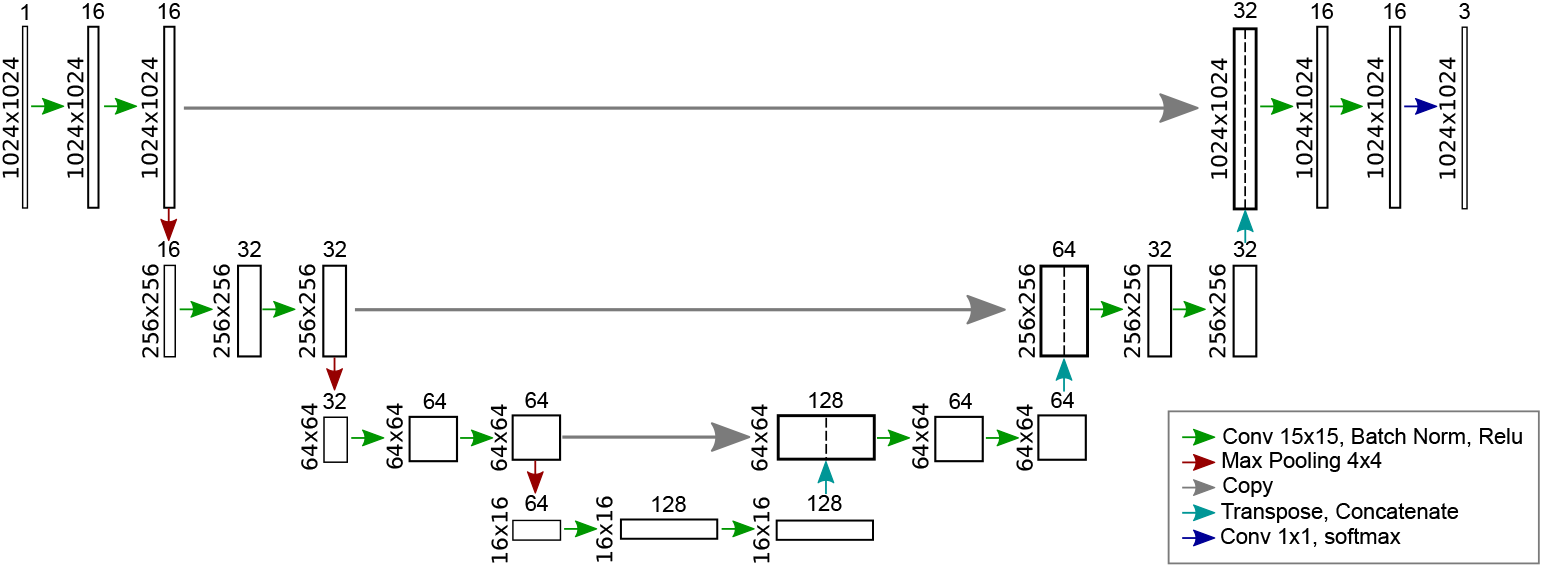
Proposed U-net model architecture for mesenchyme and neural ectoderm segmentation in DAPI-stained images.

### 2.5. Cell segmentation

U-nets have been previously used for automatic cell segmentation in various imaging modalities [15, 16, 28]. However, one of the major drawbacks identified in previous studies is the lack of annotated datasets to successfully train U-nets for the different microscopy imaging techniques available. A main contribution of this work is the establishment of two datasets (DAPI-Cells and PHH3-Cells) that facilitate the training of CNN models to segment cells in LSM images.

Owing to the limited availability of microscopy datasets, Falk et al. [16] trained and evaluated a U-net model for segmenting cells by combining images acquired using different microscopy technologies. Based on this, they developed an ImageJ-Fiji plugin that allows the re-training of this pre-trained U-net. Although the plugin facilitates transfer learning, we found experimentally that re-training of the pre-trained network had no performance advantage for segmenting DAPI-Cells and PHH3-Cells when compared to randomly initializing the model weights and training it from scratch with our data.

Thus, one U-net was trained from scratch using the DAPI-Cells dataset to automatically segment mesenchyme cells in the sample, which are known to be drivers of the change in facial morphology. For this, only the mesenchyme cell annotations were used as the positive class in the training process, and the neural ectoderm cells and background were considered the negative class. Manual hyper-parameter search was done using the ImageJ-Fiji plugin described above, optimizing the learning rate and the number of iterations. The best model was trained for 12,500 iterations with a learning rate of 1E-4, with randomly initialized weights.

After this, a second U-net was trained from scratch based on the ImageJ-Fiji plugin using the PHH3-Cells dataset, for segmenting proliferating cells in PHH3 images. Based on a manual hyper-parameter search, the best model was trained for 20,000 iterations with a learning rate of 1E-6, with randomly initialized weights.

These U-net models were compared with the Ilastik segmentation method, which is a reference tool and widely used for semiautomatic cell segmentation in biology [14]. In this tool, the user specifies the image features to be extracted and the parameters of the machine learning method, and provides microscopy datasets with corresponding cell segmentations. Due to the number of tunable variables, its effectiveness depends on the domain expertise of the user. In case of LSM, it is challenging to optimize the tool so that it generalizes and performs well on future scans due to the variability between images and the large size of them. The best results in this study were achieved using a random forest classifier integrating features based on intensity, edge, and texture. These features were computed with sigmas of 3.5, 5, 10, and 15 pixels, independently for the DAPI-Cells and PHH3-Cells sets. Pixels with a score greater than 0.3 were assigned to the positive class.

### 2.6. Full volume segmentation

Once the U-nets for segmentation of the tissues, cells, and proliferating cells are trained, full embryos can be segmented.

For cell and tissue segmentation (Fig. 2) in a new unseen LSM dataset, each image of the DAPI-stained z-stack has to be normalized and cropped to 1024 × 1024 patches so that it can be processed by the developed CNN models. Particularly for tissue segmentation, we found experimentally that an overlap of neighbouring patches is necessary to avoid discontinuities of the segmentations at the borders of the patches. Once the segmentation of cells and tissues is completed and the patches are merged, the mesenchyme segmentation is used to mask the cell segmentation so that only cells in mesenchyme remain in the segmentation.

For segmentation of proliferating cells, the images of the PHH3 volume also have to be intensity-normalized and cropped to 1024 × 1024 patches. These patches are segmented using the U-net trained with the PHH3-Cells dataset. Then, the segmented patches are merged and masked using the mesenchyme segmentation, to obtain the proliferating cells in the mesenchyme (Fig. 2).

Finally, the overall relative cell proliferation map is obtained by calculating the ratio of proliferation in a fixed 3D window centered around each voxel. For this, the number of segmented voxels corresponding to proliferating cells in mesenchyme is counted in this window and divided by the number of segmented voxels corresponding to cells in the same window.

### 2.7. Evaluation metrics

The automatic segmentations were quantitatively compared to the corresponding manual segmentations in the three test sets using the accuracy and F-score. Furthermore, a global measure for assessing the tissue segmentation is required, since it is a multi-class problem (mesenchyme, neural ectoderm, background). In this case, the macro-average F-score (Fscore_*M*_) was used, which is the average of the same measures for all classes [29].

## 3. Results

### 3.1. Tissue segmentation

The hyper-parameter search to optimize the tissue segmentation U-net CNN model was performed using Compute Canada, requesting an NVIDIA V100 Volta (32G HBM2 memory). Using the identified optimal set of parameters for segmenting the validation set (Fscore_*M*_ =0.8), a CNN model was trained using the training and validation set for segmenting the DAPI-Tissue test set using a PC with a AMD Ryzen 5 3600 3.6 GHz 6-core CPU, 32 GB RAM, and an NVIDIA GeForce RTX 2070 Super GPU.

Table 2 shows the different metrics for the automatic segmentation of mesenchyme and neural ectoderm for the validation and test sets. Each metric (F-scores, Accuracy) was computed for each image in the test and validations sets, and the global mean and standard deviation for was computed for each set. Fig. 4 shows the box-plots for the metrics on the test set.

**Table 2:** Metrics for the segmentation of mesenchyme (Mes), neural ectoderm (NE) and Background (B) in DAPI-Tissue images using the proposed U-net.

|  | **Accuracy** | **Fscore<sub>M</sub>** | **F-score Mes** | **F-score NE** | **F-score B** |
| --- | --- | --- | --- | --- | --- |
| Inter-observer (Test) | 0.971(0.031) | 0.885(0.122) | 0.849(0.196) | 0.777(0.261) | 0.916(0.146) |
| U-net (Val) | 0.900(0.08) | 0.804(0.133) | 0.783(0.209) | 0.515(0.323) | 0.974(0.015) |
| U-net (Test) | 0.889(0.093) | 0.837(0.137) | 0.799(0.203) | 0.705(0.337) | 0.931(0.141) |

**Figure 4:**
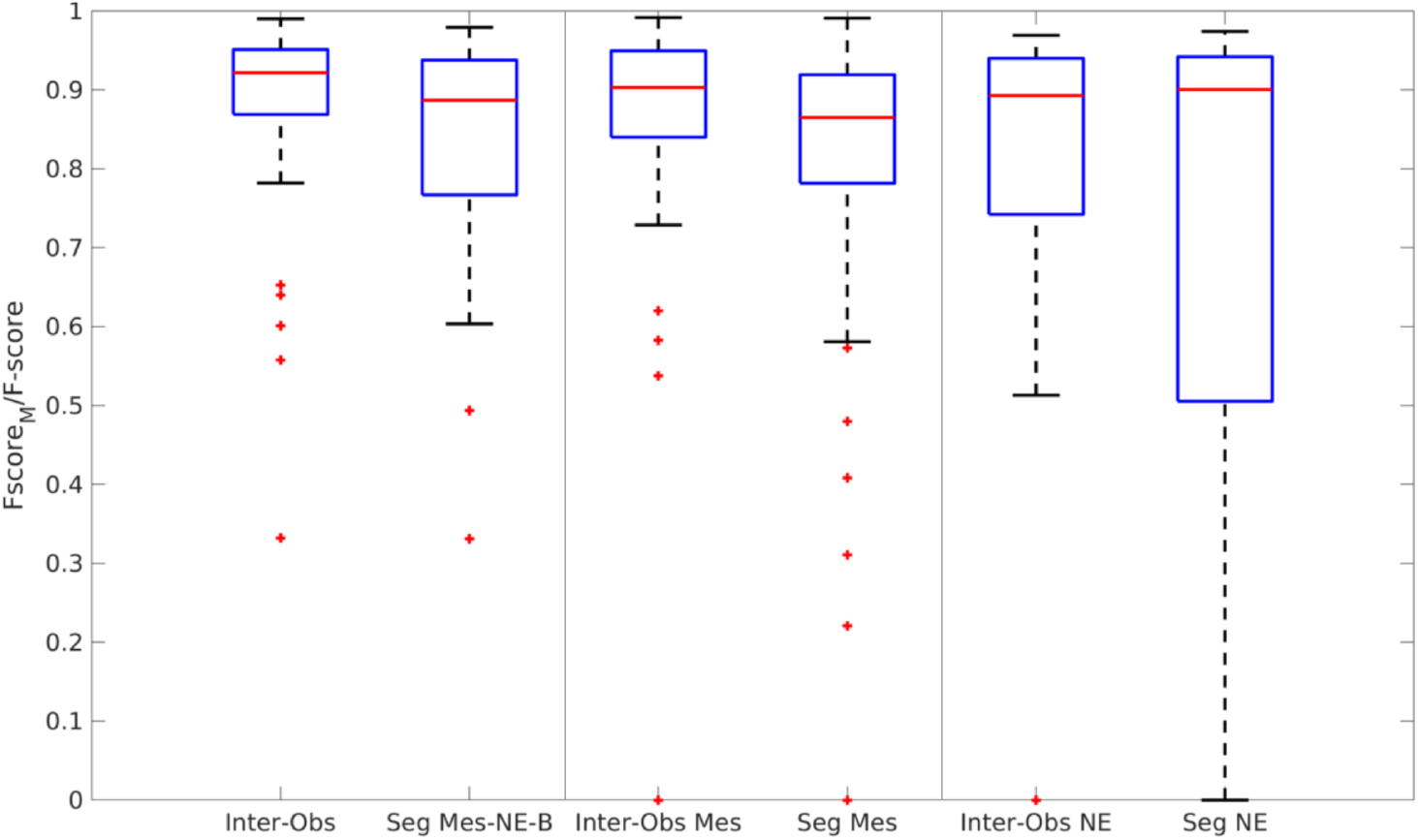
Box-plots of the overall Fscore_*M*_ and F-scores for neural ectoderm (NE) and mesenchyme (Mes) segmentation with the corresponding inter-observer agreement results for reference.

The results show that the tissue segmentation using the proposed CNN model is similar to the manual segmentations. More precisely, a high accuracy of 0.889 is reached for the test set, showing great agreement with the observers. Considering the Fscore_*M*_, a metric more appropriate for imbalanced problems [29], the value slightly decreases to 0.837. However, compared to the Fscore_*M*_ of 0.885 for the inter-observer agreement and investigating the corresponding box-plots reveals that the results of the CNN model are generally in the range of the inter-observer variability. The binary F-scores for the mesenchyme and neural ectoderm segmentations (0.799 and 0.705, respectively) demonstrate high agreement with the observers, but slightly lower than the inter-observer variability (0.849 and 0.777, respectively). The analysis of the F-score for the background classification complements the understanding of the automatic segmentation performance. In this case, a mean value of 0.931 is achieved, which is similar to the mean inter-observer agreement (0.916).

Fig. 5 shows the Bland-Altman plots for the percentual error of the segmented areas, as the segmented areas vary between datasets. The horizontal distribution around 0% suggests that the error is equally distributed indicating that no tissue is over- or under-estimated.

**Figure 5:**
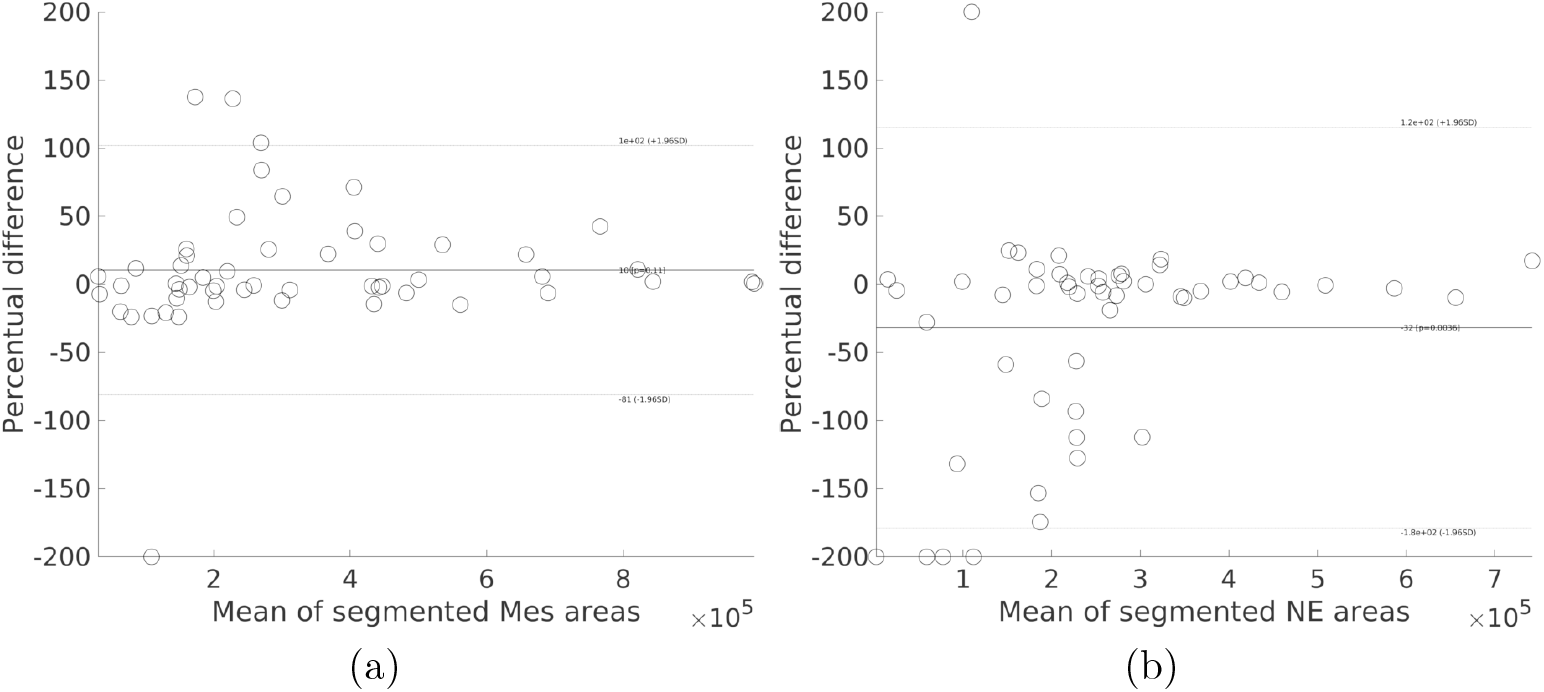
Bland-Altman plots for tissue segmentation. Areas are expressed in pixels. (a) Mesenchyme (Mes) segmentation. (b) Neural ectoderm (NE) segmentation.

Fig. 6 shows three images from the DAPI-Tissue test set with their manual segmentations and their correspondent automatic segmentation. The qualitative results are well in line with the quantitative results, showing good agreement of the automatic segmentations with the manual annotations. There are some misclassifications of mesenchyme and neural ectoderm, but the sample is properly differentiated from the background, correctly pre-serving the concavities of the embryo.

**Figure 6:**
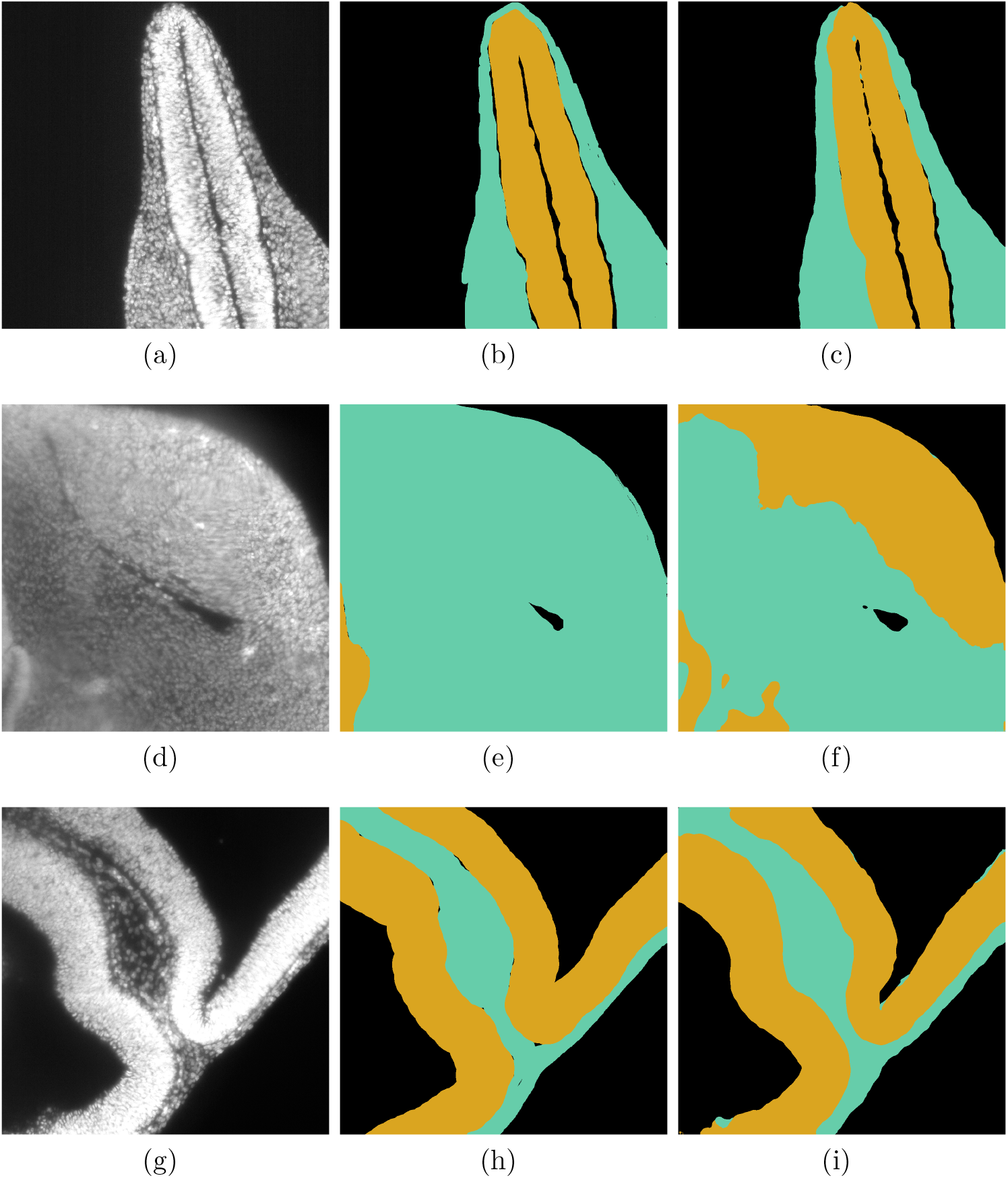
Segmentation of tissues in DAPI-stained images. (a) (d) (g) Original images, from the DAPI-Tissue test set. (b) (e) (h) Ground truth, where the mesenchyme is coloured in aquamarine and the neural ectoderm in yellow. (c) (f) (i) Segmentation result using the proposed U-net for tissue segmentation.

### 3.2. Cell segmentation

The fine-tuning of the CNN models for cell segmentation was performed on the same PC described in Section 3.1. The segmentation using Ilastik was executed on a PC with an Intel i7-8700K 3.7GHz 6-core CPU, 64 GB RAM, and a NVIDIA GeForce GTX 1080 Ti graphic card.

Table 3 shows the evaluation metrics for the automatic segmentation of mesenchyme cells in DAPI-stained images (DAPI-Cells) and proliferating cells in PHH3-stained images (PHH3-Cells) calculated for the corresponding independent test sets, including only images where the observer annotated at least one cell.

**Table 3:** Metrics for mesenchyme cell segmentation in DAPI-stained images (DAPI-Cells, test set), and proliferating cells in PHH3 images (PHH3-Cells, test set).

|  | DAPI-Cells |  | PHH3-Cells |  |
| --- | --- | --- | --- | --- |
|  | Accuracy | F-score | Accuracy | F-score |
| Inter-observer (Test) | 0.811(0.096) | 0.385(0.279) | 0.987(0.006) | 0.452(0.133) |
| Ilastik (Test) | 0.797(0.097) | 0.399(0.209) | 0.989(0.005) | 0.537(0.134) |
| U-net (Test) | 0.812(0.081) | 0.569(0.156) | 0.994(0.004) | 0.560(0.172) |

For the mesenchyme cell segmentation in DAPI images, Fig. 7 presents the boxplots for the F-scores when segmenting the test set of DAPI-Cells, and the corresponding Bland-Altman plot for the percentual error of the segmented areas. Table 3 and Fig. 7a show that the CNN model achives an F-score of 0.569, which surpasses the Ilastik results (0.399). Fig. 7b shows that the segmentation error is equally distributed around 0%, indicating no systematic bias. Qualitatively, Fig. 8 shows segmentation examples in mesenchyme regions. Here, it can be seen that the cells have blurry edges in the DAPI images. Comparing the CNN-based segmentation results with the segmentation results computed using Ilastik, it can be observed that the proposed CNN-based model produces sharper segmentations than Ilastik, which look more similar to the ground truth.

**Figure 7:**
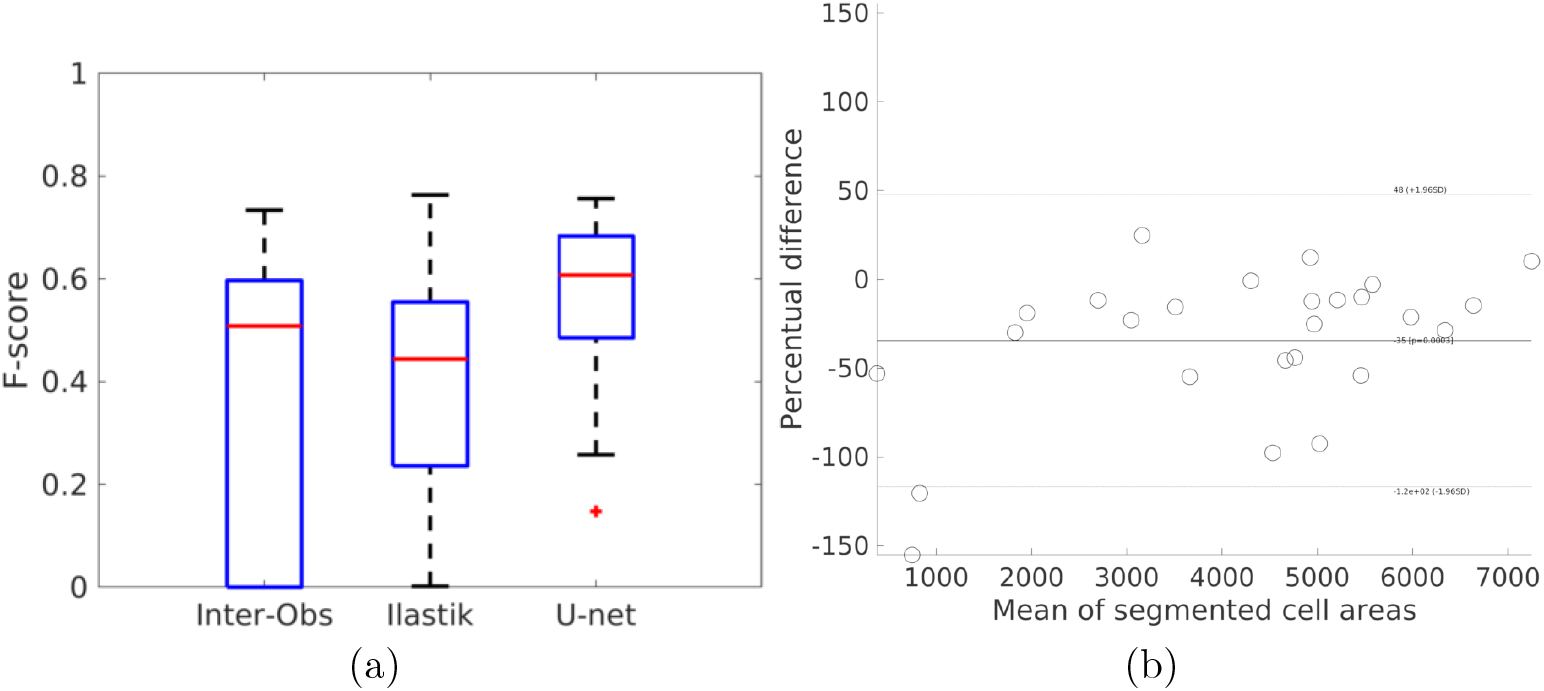
F-score for cell segmentation in mesenchyme regions in DAPI-stained images using the DAPI-Cells test set. (a) F-scores for Ilastik and U-net, compared with the inter-observer agreement. (b) Bland-Altman plot. Areas are expressed in pixels.

**Figure 8:**
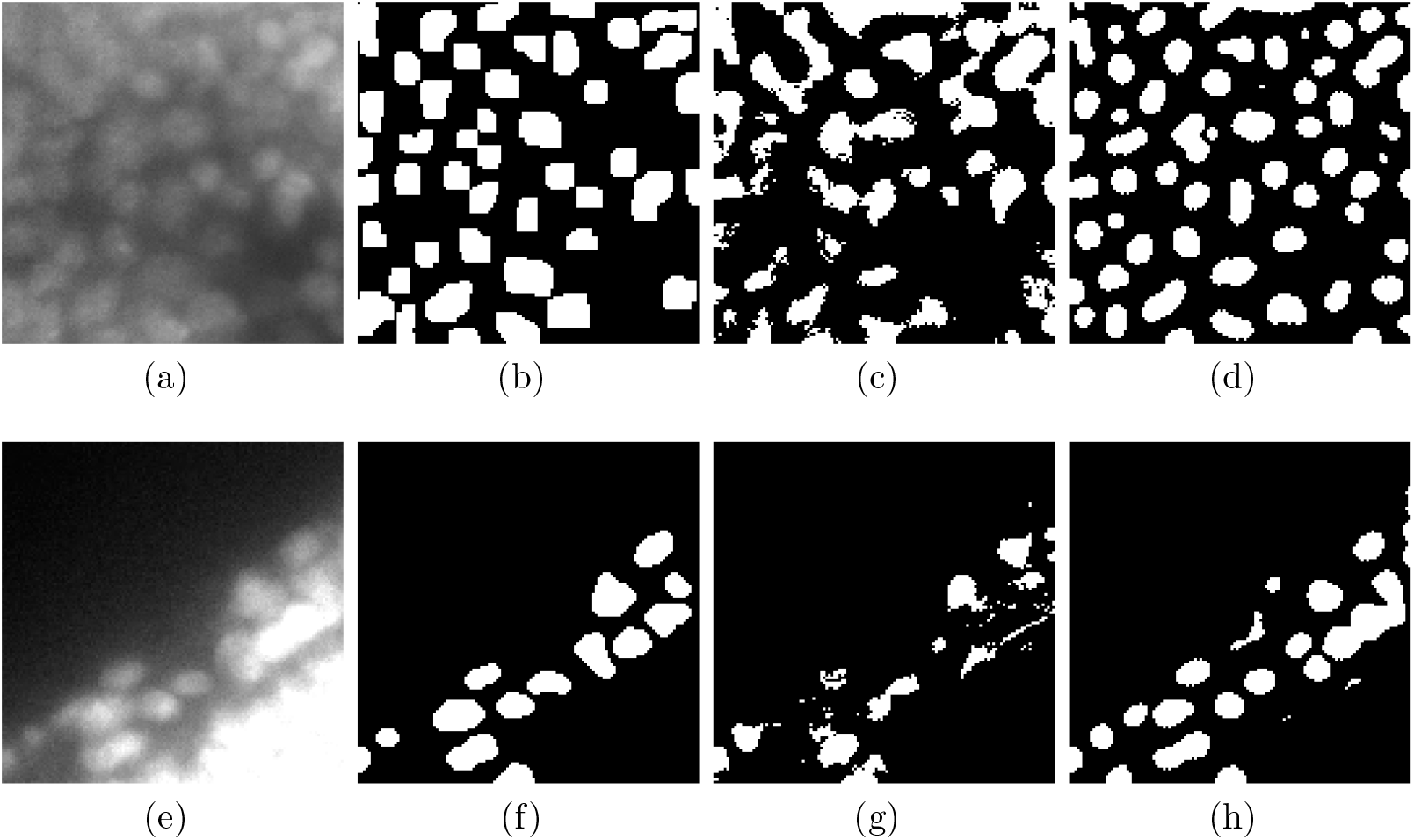
Segmentation of cells in DAPI-stained images. (a) (e) Original images from the DAPI-Cells test set. (b) (f) Ground truth in which only mesenchyme cells are segmented. (c) (g) Segmentation result using Ilastik. (d) (h) Segmentation result using U-net.

Fig. 9 shows the boxplots for the F-scores obtained for the PHH3-Cells test set and the associated Bland-Altman plot. The automatic CNN-based method achieved an F-score of 0.56 compared to the lower mean inter-observer agreement F-score of 0.45. Fig. 9b indicates a tendency towards over-estimation of the automatic CNN-based method. Fig. 10 shows examples of the segmentation of proliferating cells. Fig. 10a and 10e show that the PHH3 data presents high levels of noise and varying levels of intensities in the cells of interest. Comparing the CNN-based with Ilastik segmentations for the images, the CNN-based segmentations are affected less by noise compared to Ilastik, and are thus quantitatively slightly more accurate (F-score of 0.56 vs. 0.54).

**Figure 9:**
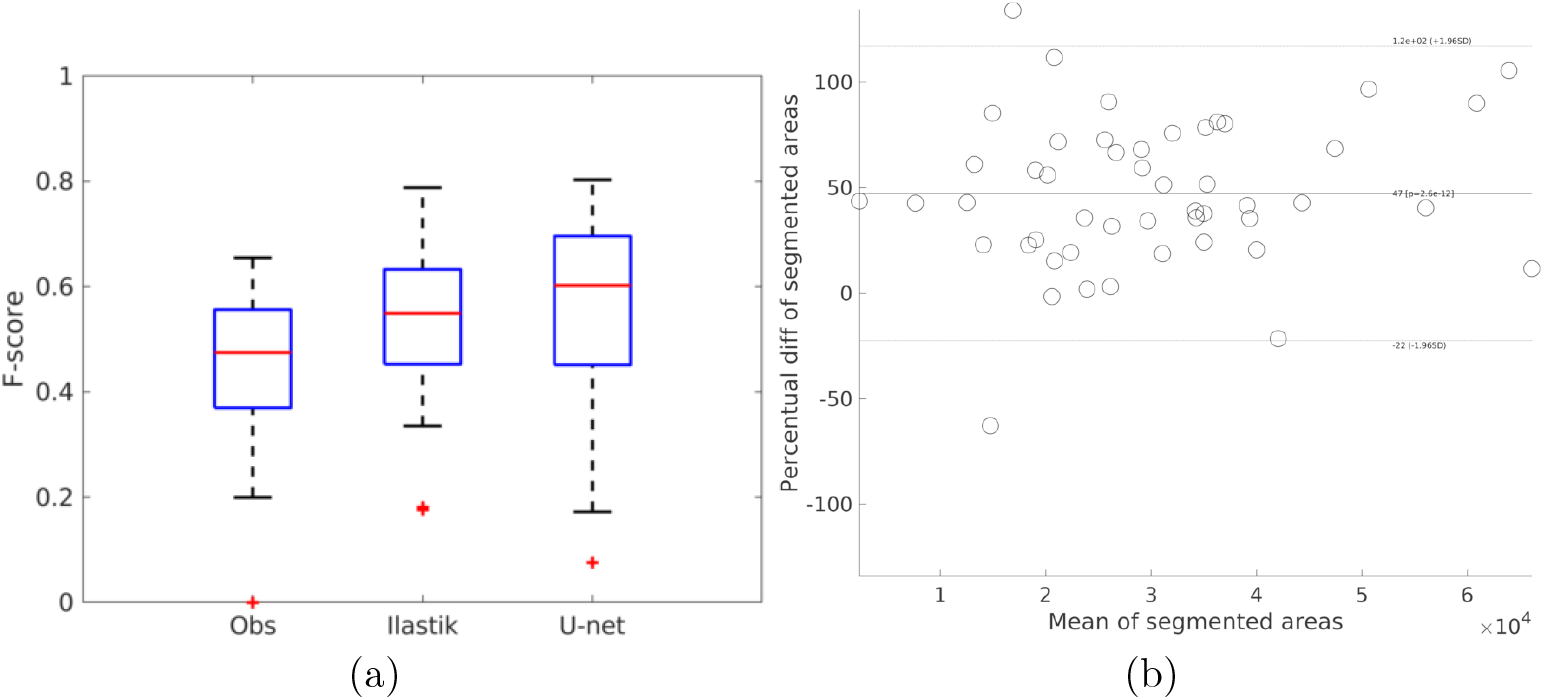
F-score for proliferating cell segmentation in PHH3 images. (a) F-scores for Ilastik and U-net compared to the inter-observer agreement. (b) Bland-Altman plot. Areas are expressed in pixels.

**Figure 10:**
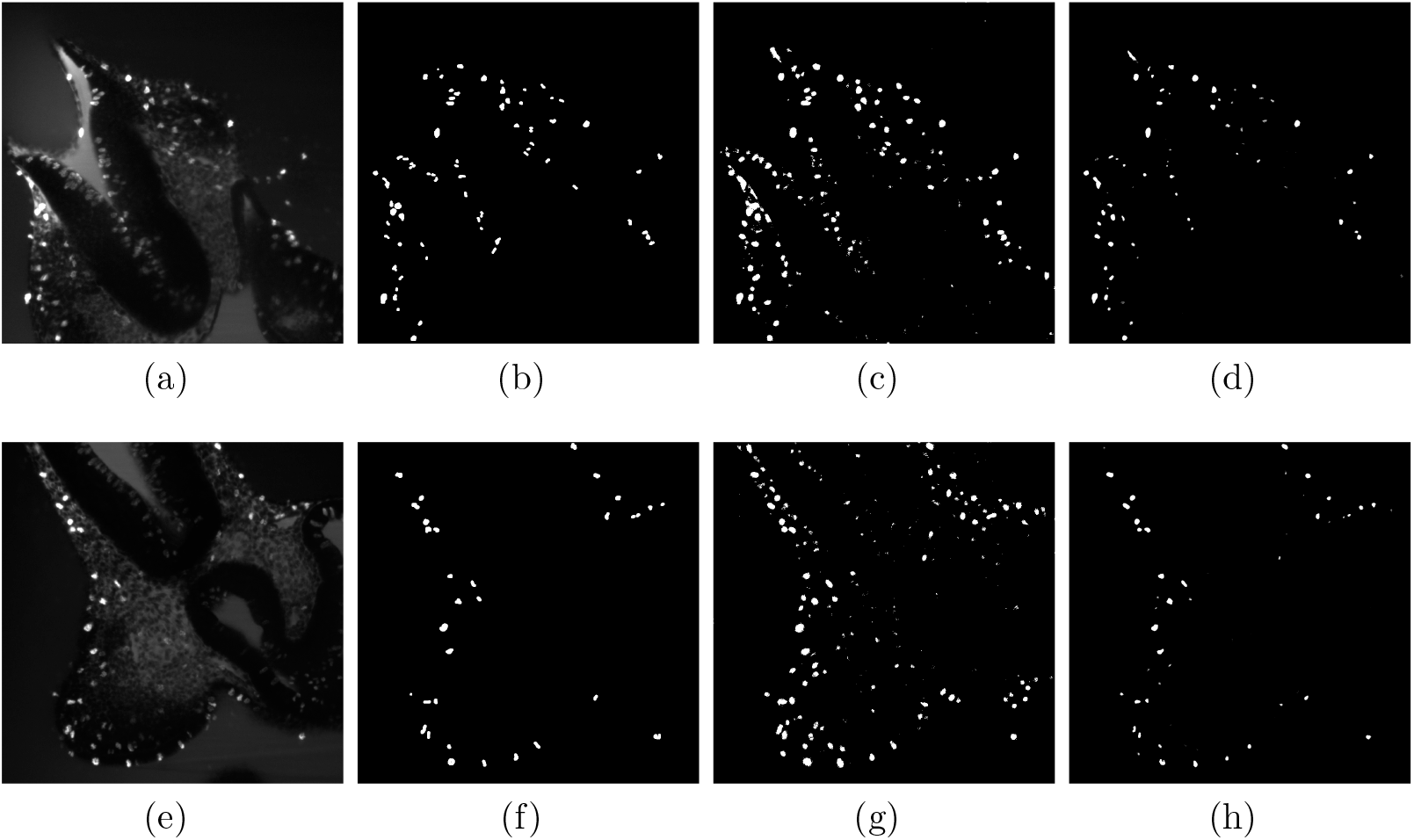
PHH3-stained image segmentation. (a) (e) Original images from the PHH3-Cells test set. (b) (f) Ground truth. (c) (g) Segmentation result using Ilastik. (d) (h) Segmentation result using U-net.

### 3.3. Workflow integration

Once the U-nets for segmentation of the tissues, cells, and proliferating cells were trained, a full E9.5 embryo head was segmented (Fig. 11) using these models. Particularly, an overlap of the patches of 100 pixels on each side was used for tissue segmentation. The resulting segmentations (tissues, cells, and proliferating cells) were downsampled to generate isotropic volumes of 297 × 297 × 407. Then, a window of 40 × 40 × 40 was used for computing the relative cell proliferation map.

**Figure 11:**
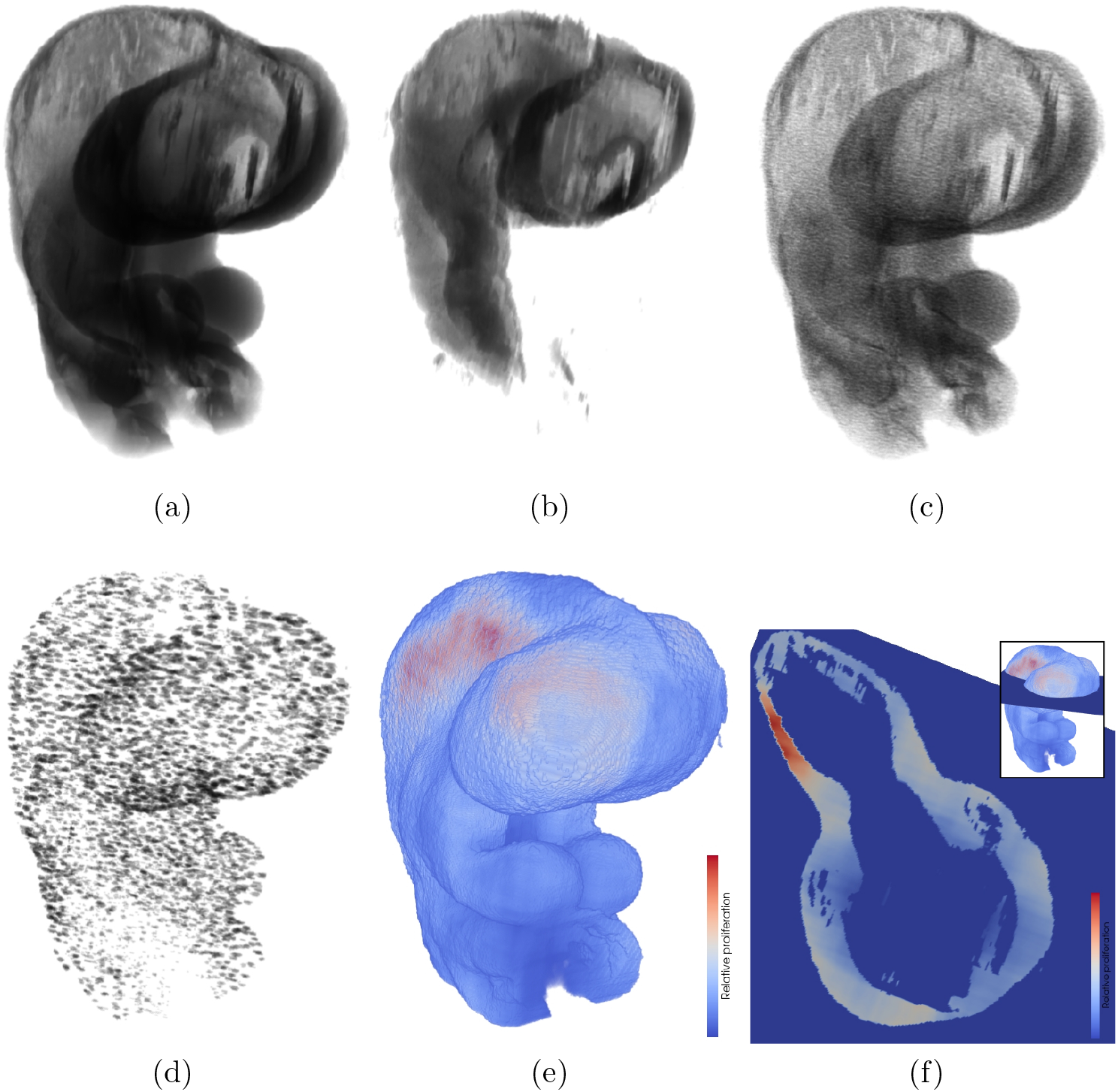
Automatically segmented E9.5 embryo. Voxels are displayed slightly transparent for visualization purposes. (a) Mesenchyme tissue segmented from DAPI images. (b) Neural ectoderm tissue segmented from DAPI images. (c) Cells in mesenchyme segmented from DAPI-stained images. (d) Proliferating cells segmented in PHH3-stained images, only in mesenchyme, masked using the tissue segmentation result. (e) Heat map of the proliferating cells, relative to all cells in the region. (f) Axial plane located in the head of the embryo, showing the relative proliferation in mesenchyme.

Qualitatively, the volume of the segmented neural ectoderm (Fig. 11b) has the expected shape for the embryonic age, and it fits inside the segmented mesenchyme (Fig. 11a), as it should. The total number of segmented cells in the scan (Fig. 11c) is higher than the total number of proliferating cells, which can be clearly seen in Fig. 11d. Using these numbers, the relative proliferation map is computed (Fig. 11e). 11f shows the relative proliferation in the mesenchyme and an empty space corresponding to the neural ectoderm.

## 4. Discussion

### 4.1. Tissue segmentation

The results suggest that the proposed CNN-based framework is able to segment the background as well as a human observer but it can can lead to some misclassifications of mesenchyme and neural ectoderm. Considering the design of the proposed CNN model, the similar validation and test results (Table 2) show that it has a good generalization capability, not over- or under-fitting the training data.

In Fig. 6, it can be seen that the embryo is well separated from the background, but the tissues are partly mislabeled. For example, in Fig. 6f, mislabelling occurs in blurry regions of the image (bottom left) and where the signal varies significantly (top right). In this case, these artifacts in the image are a result of the staining process and LSM acquisition artefacts. However, the proposed CNN-based segmentation method still leads to good results in some of these regions with less severe artefacts.

Overall, the good performance of the automatic tissue segmentation enables the use of subsequent geometric morphometrics analysis, which is a standard for developmental biology research.

### 4.2. Cell segmentation

With respect to the cell segmentation, the results show a rather low inter-observer agreement with an F-score of 0.39 for cell segmentation in DAPI-stained images and 0.45 for cell segmentation in PHH3-stained images. This low agreement has also been reported for other fluorescence microscopy datasets, and is likely related to the smaller surface area to volume ratio of cell structures compared to tissues [30] as well as blurry cell edges even where the focus is optimal. In order to assess the extent to which the quality of the scans affects the human and automatic segmentation, the image focus assessment method proposed by Yang et al. [11] was used. In DAPI-Cells test images with low defocus scores (between 1 and 5), the F-score comparing the manual segmentations was 0.49±0.22, while comparing the automatic segmentation results with the manual segmentation results led to an F-score of 0.63±0.1. For images with high defocus scores (between 6 and 11), the inter-observer F-score is 0.26±0.27, while the corresponding value for the automatic segmentation is 0.47±0.17. Thus, the loss of focus affects the manual segmentation much more than the automatic segmentation. Overall, it can be concluded that the low effectiveness of the automatic segmentation is strongly related to the quality of the training data, which includes the image quality but also the manual segmentations used.

For PHH3 segmentation, the CNN-based segmentation method achieved an F-score of 0.56, whereas the F-score of the corresponding mean inter-observer agreement was 0.45. Taking into account the overestimation of 50% observed in Fig. 9b, it can be argued that the automatic segmentation does not perform optimal due to non-specific fluorescent signals. These artifacts in the image resemble proliferating cells, which lead to falsely segmented voxels in the manual as well as automatic segmentations.

However, it needs to be highlighted that the evaluation metrics of the proposed CNN-based method for segmentation of cells are better than the corresponding metrics for the Ilastik segmentation. Qualitatively, the proposed CNN-based segmentation method produces better defined cells. The improved sensitivity of the CNN-based method compared with the Ilastik results is likely a consequence of taking high level textural features into account, which are computed in deeper CNN layers. These features are not available or computed in the Ilastik method, even when it is configured by users trained in image processing.

### 4.3. Workflow integration

Fig. 11 shows each segmentation result as generated by the proposed workflow in a full E9.5 embryo head. All segmentations fulfill the expected properties. However, some unexpected sagittal asymmetries are present. They can be seen in the resulting mesenchyme shape and in the distribution of proliferating cells in Fig. 11f. These asymmetries could be the result of variance in the staining of the embryo, in both DAPI and PHH3, its pose during the scanning, the laser penetration, and errors in the automatic segmentation methods. Asymmetries could affect further shape and morphological analysis. As a sagittal symmetry of their morphology and proliferation can be assumed in wild-type mice, this assymetry can be corrected in the segmentations using standard post-processing methods such as affine deformations based on an embryo atlas or predefined landmarks.

Generally, the overall results show that the CNN models can segment the structures of interest accurately and in the range of human observers. Thus, this automatic segmentation workflow can be efficiently used for quantitative analysis of LSM images, oriented to developmental biology studies. However, the inter-observer metrics show that there is room for improvement with respect to the quality of the image acquisition.

## 5. Conclusion

The aim of the present work was to implement and evaluate an automatic framework for tissue and cell segmentation in LSM images. Proper tissue and cell segmentation is important for the quantification and modeling of how perturbations to cellular dynamics result in congenital anomalies. Such analyses are based on data that rely heavily on extraction of variables such as cell number, size, or density from noisy volumetric image data. Thus, a database of LSM images was established with corresponding manual segmentation of proliferating cells, tissues, and total cells. One CNN model was trained for each segmentation problem, and the quantitative evaluation suggests that all three models lead to segmentation results within the range of the inter-observer agreement. The source code, software, and annotated datasets are publicly available at https://github.com/lucaslovercio/LSMprocessing. The methods developed in this work are integral to the larger goal of improving the understanding of development and morphogenesis and how perturbations to development result in diseases.

## Acknowledgements

The work was supported by NIH NIDCR R01-DE019638 to R. M. and B. H., NSERC Discovery to B. H., and CIHR Foundation grant to B. H. and R. M. Funding sources had no input in experimental design or interpretation.

LL is supported by an Eyes High postdoctoral fellowship (University of Calgary), RG by a CIHR fellowship, MM by a Cumming School of Medicine and McCaig Institute postdoctoral fellowship, MVG by an Alberta Children’s Hospital Research Institute Postdoctoral Fellowship, and NDF by the Canada Research Chairs Program.

The financial support of these institutions is greatly appreciated.

This research was enabled in part by support provided by WestGrid and Compute Canada (www.computecanada.ca).

The authors thank Divam Gupta from Carnegie Mellon University (www.divamgupta.com) for guidance in semantic segmentation.

